# Blockages to biodiversity data access for conservation and sustainability management

**DOI:** 10.64898/2026.06.13.732021

**Authors:** P.J. Stephenson, Kerrigan Marie Machado Unter, Judith L. Walls, Jorge Armando Amador Moncada, Louisa Sawyerr, María Cecilia Londoño Murcia, Yaa Ntiamoa-Baidu, Luca Fumagalli

## Abstract

Governments, civil society organizations and businesses often lack the biodiversity data they need for decision-making and adaptive management, impacting their planning, reporting and performance. We explored the biodiversity data needs of such actors in Colombia, Ghana and Switzerland to identify factors affecting data availability and use. Responses to questionnaire surveys showed that the data types with the biggest gaps between user needs and access were progress on conservation or sustainability actions, species populations, habitat state and ecological risk. The most frequent data blockages related to inadequate resources and organizational capacity. Obstacles significantly associated with a lack of primary data included an absence of organizational biodiversity goals and monitoring systems. Problems accessing habitat quality and species abundance data were associated with data collection methods being unknown or unavailable. Businesses were more likely than other groups to need data on threats, perhaps reflecting the increasing importance of environmental risk to the corporate sector. Businesses are less likely to collect primary data or use secondary data and are significantly more likely to be unclear on what biodiversity indicators to use. Non-business organizations are significantly more likely to be unable to access data because of a lack of funding for data collection, analysis, and use. Our results highlight the need for stakeholders across sectors to work together to find common solutions to build and invest in monitoring capacity that unblocks the flow of biodiversity data.

## INTRODUCTION

Biodiversity is declining (Secretariat of the Convention on Biological Diversity 2020, WWF 2024). Impactful action to reverse current trends requires effective, results-based decision-making and adaptive management. To that end, many stakeholders need data on the state of species and habitats, the pressures they face, the benefits accrued from ecosystem services, and their management and policy responses. Governments need data, for example, to develop environmental legislation and policies, manage resources across industries (e.g., agriculture, fisheries, mining), and deliver multilateral environmental agreements (Stephenson et al. 2017). Businesses need biodiversity information to meet sustainability targets, monitor and report their environmental impacts, and manage risk (Unter et al. 2023, 2024, Testa et al. 2025, IPBES 2026). Conservation non-governmental organizations (NGOs) need data to prioritize actions, monitor outcomes and impacts, and apply adaptive management (CMP 2020). Most actors also need data to demonstrate contributions to global goals and policy processes, such as the Sustainable Development Goals (United Nations 2023) and the Kunming-Montreal Global Biodiversity Framework (Convention on Biological Diversity 2022).

However, biodiversity data are frequently scattered, fragmented, of poor quality, and rarely available in the right format at the right time (Kissling et al. 2018, Stephenson et al., 2022). Consequently, government reporting on biodiversity often lacks data (Walpole et al 2009, Bubb 2013, Convention on Wetlands, 2025) and few companies report on biodiversity (Overbeek et al. 2013, Adler et al. 2018, Azizi et al. 2025). Conservation NGOs also struggle to collect and use data to monitor their impacts on biodiversity (McKinnon et al. 2015, Moreno et al. 2023). These gaps ultimately jeopardize the performance and impact of conservation and sustainability policies and actions.

A number of challenges have been identified that prevent the use of biodiversity data in decision-making. These include a lack of capacity and tools for identifying indicators and collecting, analysing and interpreting data (Stephenson et al. 2017, 2022, Addison et al. 2020, Hochkirch et al. 2020). Biodiversity monitoring schemes and databases have taxonomic and geographic biases and data access limitations (Troudet et al. 2017, Tydecks et al. 2018, Moussy et al. 2022, Kemp et al. in press), and many institutions fail to follow data management best practices (Wilkinson et al. 2016, Augustine et al. 2024). Variability in the spatial and temporal resolution of data, a lack of willingness to share information, and the failure to link risks and dependencies to actions, also affect governments and businesses (Bansal & Knox-Hayes 2013, Whiteman et al. 2013, Stephenson et al. 2022, Lahneman et al. 2025). For example, in the tourism sector in Iceland, many relevant environmental indicators—including indicators on endangered species, protection of areas, soil erosion, and water quality—cannot be measured due to lack of data, disaggregation issues, and the omission of metrics on key pressures (Saviolidis et al. 2021). Therefore, many stakeholders struggle to identify appropriate indicators for monitoring biodiversity, sources of existing data they can use, and the relevant monitoring tools for collecting their own data (Boiral & Heras-Saizarbitoria 2017, Addison et al. 2020, Stephenson & Stengel 2020).

How can these blockages be addressed and data made more freely available to inform decision-making and enhance conservation impact and environmental sustainability? We brought together experts from multiple disciplines to explore biodiversity data user needs in three sample countries on three continents: Colombia, Ghana and Switzerland. We conducted an online questionnaire survey of key data users to identify the reasons behind blockages to data access and used the results to identify potential solutions. This was therefore one of the first studies to compare biodiversity data access challenges with the data needs of different users.

Our main research questions were: What are the biodiversity data needs of international organizations, governments, civil society organizations and businesses? What factors curtail biodiversity monitoring and data access and thereby block the flow of information for decision-making?

## MATERIALS AND METHODS

### Methods

We conducted a questionnaire survey of a range of stakeholders in the three target countries. Participants were chosen from people identified in the literature review from each main stakeholder group: government, international organization, civil society organization and business. Colombia, Ghana and Switzerland were selected based on existing research collaborations and the fact they offered a representative sample of countries with diverse institutional mechanisms as well as varied biological and socio-economic contexts across which to identify potential global trends in data needs and access.

Questionnaire surveys are an often-used tool in conservation biology for assessing people’s opinions on key biodiversity topics (Chen et al. 2019, Danovaro et al. 2020, Mair et al. 2021). Such surveys are a form of expert elicitation, a tool used to develop estimates of unknown or uncertain quantities based on careful assessment of the knowledge and beliefs of experts about those quantities (Morgan 2014) and used especially when data are sparse or lacking (Meyer & Booker 2001).

Surveys were conducted online using relevant platforms to create, host, and administer them (Google Forms in Colombia, KoboToolbox in Ghana, Sawtooth Software in Switzerland). We also employed snowball sampling by encouraging potential participants to forward the survey to others that they thought would have knowledge of our research topic (Heckathorn, 1997, Goodman 2011). This increased our sample size for later statistical analysis. We split the survey into two different types of surveys: the full survey and a shorter business survey where questions were condensed, simplified, or shortened. We did this to encourage the business practitioners to complete the survey, as employees with managerial or supervisory responsibilities tend to have lower response rates (Anseel et al. 2010). For example, even when providing a shorter survey as an option, the response rate in Switzerland for business practitioners was approximately 18% compared to the longer survey sent to other participants which had a response rate of 35%. In Colombia and Ghana online surveys were supplemented with face-to-face key informant interviews to enhance sample size; in these cases, the same questions were asked as in the online surveys. In all surveys, we included a question about the country to act as a check to confirm that participants were from the correct country and to ensure snowball sampling did not introduce participants outside of our countries of focus. Each survey began by informing participants of the purpose of the study, that their personal data and responses would be kept confidential, and that information from the survey would only be reported in aggregate. The surveys were conducted in 2023, from 1 May to 30 August in Colombia, from 3 March to 7 August in Ghana and from 21 March to 11 May in Switzerland.

### Measures

The surveys explored the following measures

#### Data needs and access

We measured the data needs and access for the following categories of data: Species populations; Species demographics; Habitat state; Pressures (or threats to biodiversity); Natural capital dependency risks; Ecological risk score; Availability of ecosystem services; Progress on conservation or sustainability actions, policies and other responses. We measured data needs and access through several Likert scales in our survey. For the survey sent to non-business organizations, we had a finer breakdown of types of indicators for each data category for respondents to rate their need for and access to. See Table 1 for the full breakdown. We listed each type of data with a Likert scale. For data needs, we asked respondents to rate their agreement or disagreement with the statement: “I need data on this indicator type” (1 = Strongly disagree, 2 = Disagree, 3 = Neither agree nor disagree, 4 = Agree, 5 = Strongly Agree). For data access, we asked respondents to rate their agreement or disagreement with the statement: “I can access adequate data on this indicator type to meet my needs” (0 = Not applicable, 1 = Strongly disagree, 2 = Disagree, 3 = Neither agree nor disagree, 4 = Agree, 5 = Strongly Agree).

**Table 1.** Biodiversity data blockages by country.

| Blockage | Colombia<br>(n=47) | Ghana<br>(n=40) | Switzerland<br>(n=29) | All<br>(n=116) | Rank |
| --- | --- | --- | --- | --- | --- |
| *Lack of ability or goals* |  |  |  |  |  |
| Lack of understanding of the importance of monitoring in my organization | 25.5% | 25.0% | 20.7% | 24.1% | 17 |
| Biodiversity work including monitoring is of low priority in my organization | 29.8% | 30.0% | 20.7% | 27.6% | 15 |
| Lack of measurable biodiversity goals in my organization | 31.9% | 27.5% | 37.9% | 31.9% | 13 |
| No monitoring and evaluation systems in place | 42.6% | 30.0% | 58.6% | 42.2% | 5= |
| Lack of consensus about what to monitor | 38.3% | 30.0% | 37.9% | 35.3% | 12 |
| Unclear which indicators to use | 44.7% | 30.0% | 55.2% | 42.2% | 5= |
| Lack of technical capacity for data collection, analysis, and use | 25.5% | 37.5% | 44.8% | 38.8% | 8= |
| Lack of funding for data collection, analysis, and use | 55.3% | 70.0% | 51.7% | 59.5% | 1 |
| Lack of management willingness to collect and use data | 27.7% | 32.5% | 20.7% | 29.3% | 14 |
| *Data unavailable* |  |  |  |  |  |
| Cannot access from other sources | 83.0% | 42.5% | 13.8% | 51.7% | 2 |
| Relevant data collection methods unknown or unavailable | 78.7% | 37.5% | 10.3% | 47.4% | 3 |
| *Biases in data* |  |  |  |  |  |
| Data available have taxonomic biases or gaps | 44.7% | 40.0% | 24.1% | 37.9% | 10 |
| Data available have geographic biases or gaps | 61.7% | 30.0% | 31.0% | 43.1% | 4 |
| *Unusable data* |  |  |  |  |  |
| Poor quality data | 27.7% | 30.0% | 17.2% | 25.9% | 16 |
| Inconsistent data (e.g., collected by different consultants in different ways) | 55.3% | 40.0% | 13.8% | 39.7% | 7 |
| Data in wrong scale or format | 53.2% | 37.5% | 6.9% | 36.2% | 11 |
| Data not timely or up-to-date | 63.8% | 32.5% | 6.9% | 38.8% | 8= |

#### Factors causing data blockages

Based on data blockages identified through the literature review, we measured the following types of blockages: Lack of understanding of the importance of monitoring in the organization; Biodiversity work including monitoring is of low priority in the organization; Lack of measurable biodiversity goals in the organization; No monitoring and evaluation systems in place; Lack of consensus about what to monitor; Unclear which indicators to use; Lack of technical capacity for data collection, analysis and use; Lack of funding for data collection, analysis and use; Lack of management willingness to collect and use data; Uneven geographical distribution of data. We listed each potential blockage with a Likert scale. We asked respondents to rate their agreement or disagreement with the statement: “This is a reason why I cannot access the data I need” (1 = Strongly disagree, 2 = Disagree, 3 = Neither agree nor disagree, 4 = Agree, 5 = Strongly Agree).

#### Primary data collection

We asked respondents if they collected primary data (0 = No, 1 = Yes). If they answered yes, we then asked them if they collected each of the following categories of data: Species populations; Species demographics; Habitat state; Pressures (or threats to biodiversity); Natural capital dependency risks; Ecological risk score; Availability of ecosystem services; Progress on conservation or sustainability actions, policies and other responses. We measured each category with a dummy variable (0 = No, 1 = Yes). We then asked additional questions about their primary data including whether the data were digitised (0 = No, 1 = Yes), and what data formats the data were stored in. The data formats included spatial data, multimedia data, plain or text data, and other data formats. We measured each data format with a dummy variable (0 = No, 1 = Yes).

#### Secondary data usage

We asked respondents if they used secondary data (0 = No, 1 = Yes).

#### Organization variables

We asked questions about the organization to control for organizational level variables. First, we asked respondents for the type of organization they worked for. The options for type of organization included government department; international organization; university or other research institution; civil society organization (CSO), NGO, foundation, charity, or other non-for-profit or community group; private company; public company; financial institution; third party certifier or intermediary; industry, trade, or business association; and other organization. We then created a dummy variable to measure if the organization was a business (0 = No, 1 = Yes). Second, for the business survey, we asked respondents the size of the organization (1 = 1 - 249 employees, 2 = 250 - 499 employees, 3 = 500 - 999 employees, 4 = 1,000 - 4,999 employees, 5 = 5,000 - 9,999 employees, 6 = More than 10,000 employees). Third, for the business survey, we also asked respondents what industry their organization is in (Sector 1 (e.g., raw materials, agriculture, logging, fishing, forestry, mining, etc), 2 (e.g., manufacturing, construction, industry and energy, etc.), and/or 3 (e.g., services, financial, administrative, transportation, distribution, wholesaling and retailing, food and beverage, entertainment, IT and information, etc.)). We then created dummy variables for each sector (0 = No, 1 = Yes).

#### Demographic variables

We asked questions about the respondent to control for individual variables. We asked for both the respondents’ gender (1 = Male, 2 = Female, 3 = Prefer not to specify) and age (1 = 18 – 24, 2 = 25 – 34, 3 = 35 – 44, 4 = 45 – 54, 5 = 55 – 64, 6 = 65 +).

### Data analysis

All statistical analyses were completed in Stata 16. We used descriptive statistics to examine the most prominent data needs, most common data accessed, and the prominence of data blockages for each country (see Figure 1 and Table 1). We then aimed to compare differences across businesses versus other organizations (e.g., NGOs, governments, etc.). We used t-tests to test for the differences in the means across the key variables of interest for the survey responses from individuals in businesses versus the other organizations (see Table 2). We used a t-test because it is best for small sample sizes. Before conducting the t-tests, we tested if the variances across the key variables of interest were equal or unequal in businesses versus the other organizations. If the variances were unequal, we then adjusted the t-test to account for unequal variances across the two groups.

**Figure 1.**
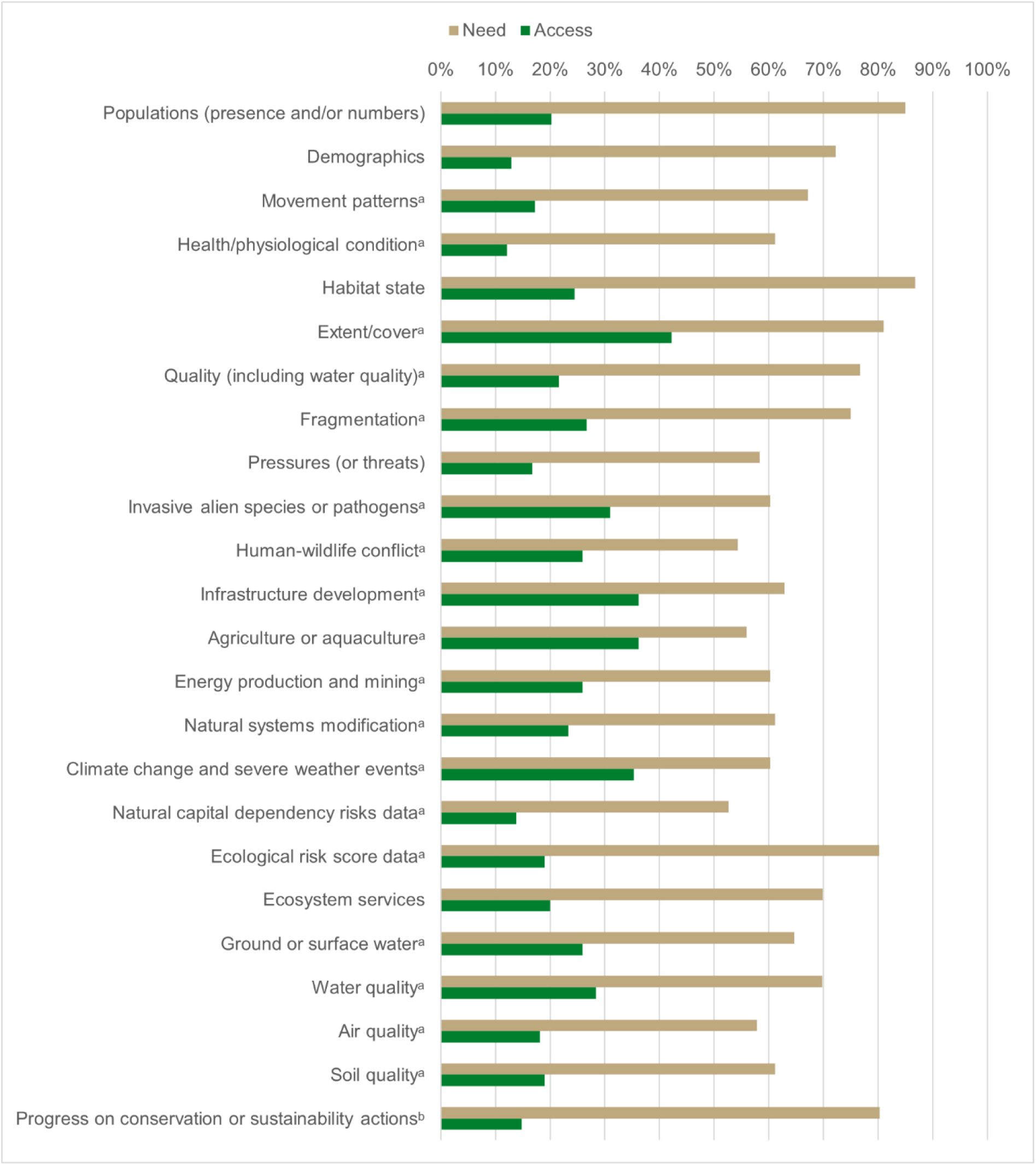
Biodiversity data needs and access for all stakeholders across all three countries. Notes: ^a^n = 116. Other variables have less observations due to incomplete survey responses; ^b^Data for Ghana not available. % = percent of respondents who replied to the question with “Agree” or “Strongly Agree”.

**Table 2.**
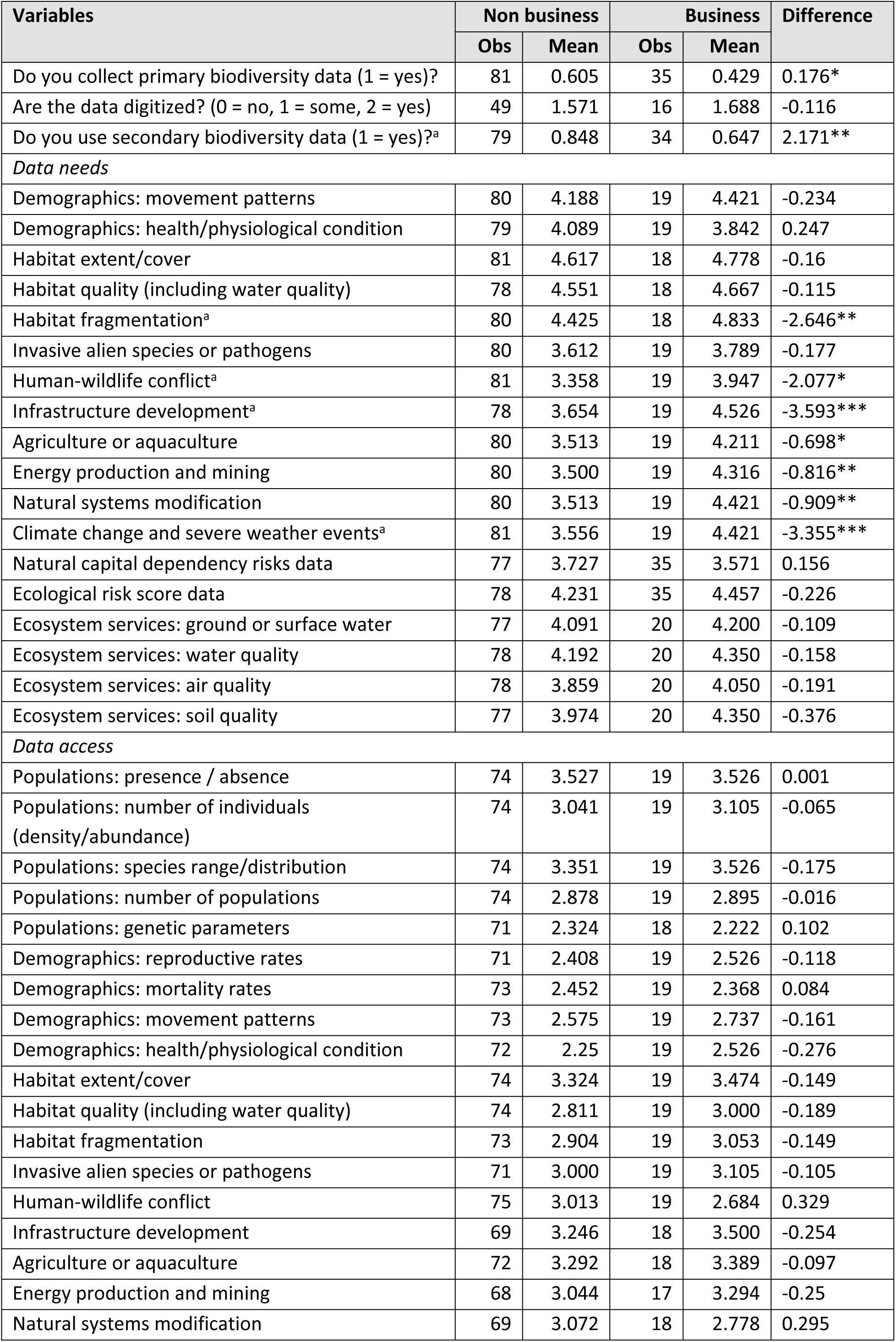
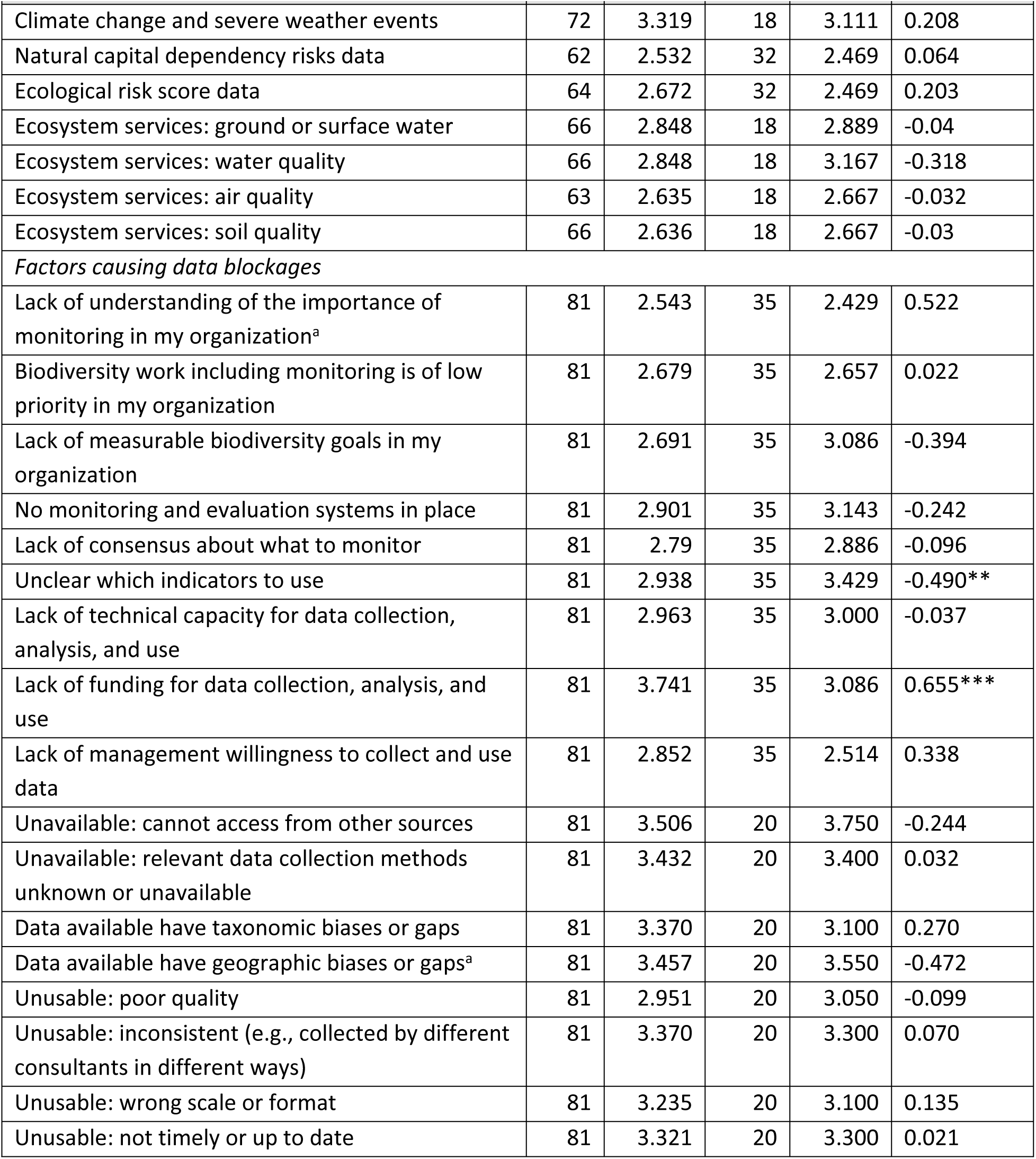
Differences in means: comparing businesses with other types of organizations. Note: ^a^Adjusted for unequal variances. *** p<0.01, ** p<0.05, * p<0.1.

Finally, we tested the relationships between variables using generalized linear models (GLMs) with robust standard errors. We controlled for respondent gender, age, and country in these models and for whether the organization was a business or not. We controlled for individual level variables such as age and gender to limit confounds that may arise from the individual filling out the survey, so that we can isolate the effect of the blockages on data collection or access. For models where the outcome was binary (e.g., whether the organization collects primary data or not), we specified a logit model.

## RESULTS

In total we received 116 completed surveys across three countries, 47 (40.5%) from Colombia (15 from businesses; 32 from other types of organization), 40 (34.5%) from Ghana (4;36) and 29 (25%) from Switzerland (16;13). Eighty-one responses were from organizations that included NGOs, governments, universities and foundations, and 35 responses were from businesses.

Sixty-four respondents indicated that their organization collects primary data on biodiversity, and 89 use secondary biodiversity data. Table 1 and Figure 1 summarise the descriptive results for variables that have data for all three countries. The types of data most needed by respondents are those to measure habitat state (especially habitat extent or cover) (86.8%) and species populations (presence and/or numbers) (85.0%), as well as progress on conservation or sustainability actions (80.3%) and ecological risk (80.2%). Data most accessible to respondents were habitat extent (42.2%) and pressures around infrastructure (36.2%), agriculture/aquaculture (36.2%) and climate change and severe weather (35.3%). Data with the biggest gaps between user needs and access (i.e., the difference between access and needs) were progress on conservation actions (65.5%), species populations (64.8%), habitat state (62.3%) and ecological risk (61.2%) (Fig. 1).

We found that data needs and access to data vary between data user groups and between countries. Data users, from government agencies to NGOs to businesses, face a diverse and complex array of blockages that prevent them accessing all the high-quality information they need in the right format at the right time to make informed decisions on the management or conservation of natural resources. For example, on average, 87% of data users require information on the state of natural habitats, yet only 25% can access such data.

Many businesses focus more on risk data and tend to use secondary data more than other user groups. For example, in Switzerland there are more issues blocking biodiversity data use for businesses than for other types of organization. Data users in Ghana most need species population data; habitat data are more important to users in Colombia; Swiss data users most need pressure data.

The six most frequently cited data blockages across all countries (Table 1) were: lack of funding for data collection, analysis, and use; cannot access data from other sources; relevant data collection methods unknown or unavailable; data available have geographic biases or gaps; no monitoring and evaluation systems are in place; unclear which indicators to use.

Comparing businesses with non-business data users (Table 2), we found that businesses on average are less likely to collect primary data or use secondary data. Businesses on average are also more likely to need data on habitat fragmentation, human-wildlife conflict, infrastructure development, agriculture and aquaculture, energy production and mining, natural systems modification, and climate change and severe weather events than other types of organizations. Businesses are significantly more likely to be unable to access data because they are unclear on which indicators to use, whereas non-business organizations are significantly more likely to be unable to access data because of a lack of funding for data collection, analysis, and use.

The GLMs (Table 3) identified some relationships between variables. In particular, the following blockages were significantly related to organizations being less likely to collect primary data: biodiversity work including monitoring is of low priority in the organization (β=-0.61, p<0.05); lack of measurable biodiversity goals in the organization (β=-0.45, p<0.05); no monitoring and evaluation systems in place (β=-0.56, p<0.05).

**Table 3.**
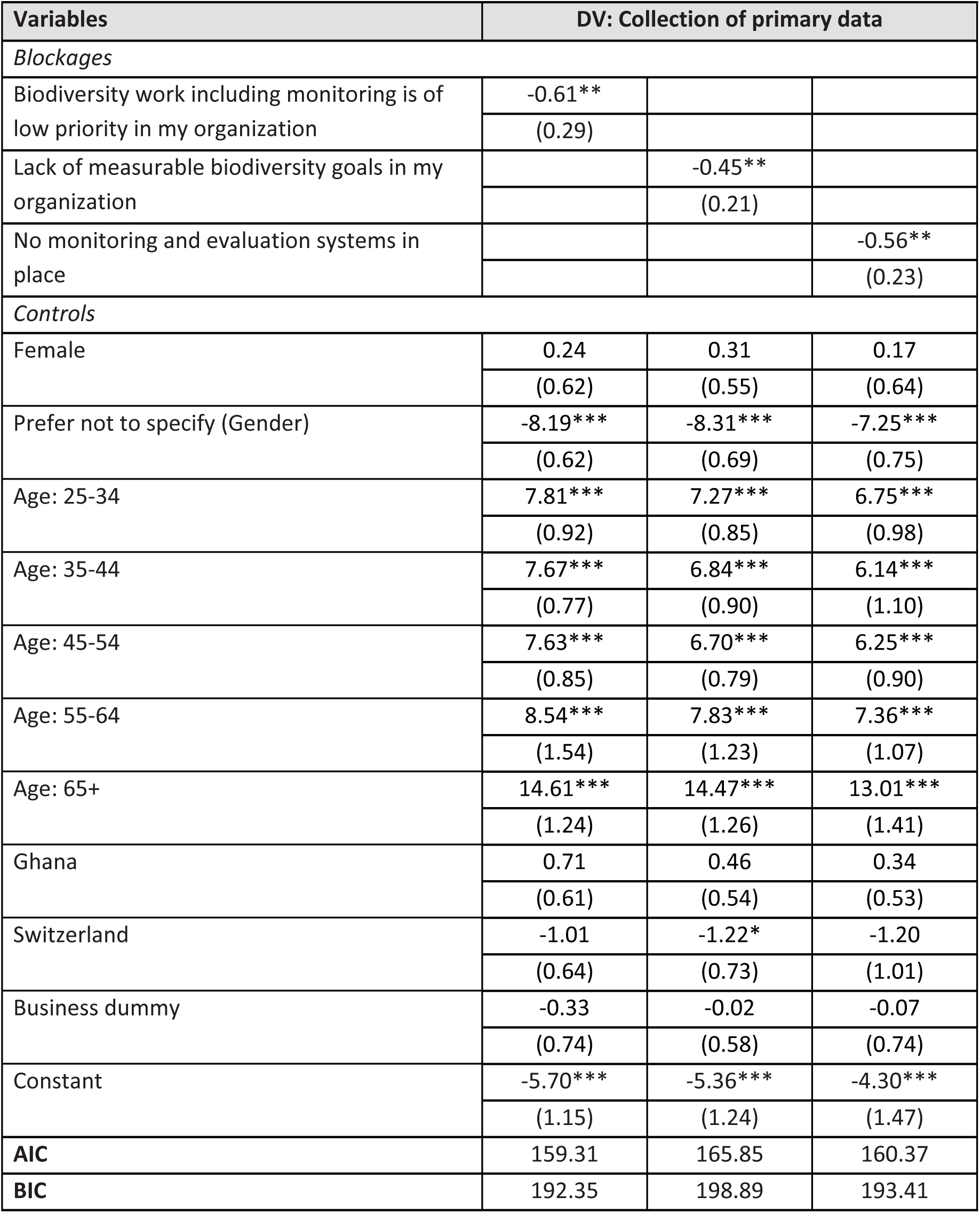
Generalized linear models for blockages against likelihood to collect primary biodiversity data. Note: n = 116. Logit model specified. DV = dependent variable. Robust standard errors in parentheses. Models where blockages are p<0.05 are presented here; full models available in Supplemental Materials. *** p<0.01, ** p<0.05, * p<0.1

| Variables | DV: Collection of primary data |  |  |
| --- | --- | --- | --- |
| Blockages |  |  |  |
| Biodiversity work including monitoring is of low priority in my organization | -0.61** |  |  |
|  | (0.29) |  |  |
| Lack of measurable biodiversity goals in my organization |  | -0.45** |  |
|  |  | (0.21) |  |
| No monitoring and evaluation systems in place |  |  | -0.56** |
|  |  |  | (0.23) |
| Controls |  |  |  |
| Female | 0.24 | 0.31 | 0.17 |
|  | (0.62) | (0.55) | (0.64) |
| Prefer not to specify (Gender) | -8.19*** | -8.31*** | -7.25*** |
|  | (0.62) | (0.69) | (0.75) |
| Age: 25-34 | 7.81*** | 7.27*** | 6.75*** |
|  | (0.92) | (0.85) | (0.98) |
| Age: 35-44 | 7.67*** | 6.84*** | 6.14*** |
|  | (0.77) | (0.90) | (1.10) |
| Age: 45-54 | 7.63*** | 6.70*** | 6.25*** |
|  | (0.85) | (0.79) | (0.90) |
| Age: 55-64 | 8.54*** | 7.83*** | 7.36*** |
|  | (1.54) | (1.23) | (1.07) |
| Age: 65+ | 14.61*** | 14.47*** | 13.01*** |
|  | (1.24) | (1.26) | (1.41) |
| Ghana | 0.71 | 0.46 | 0.34 |
|  | (0.61) | (0.54) | (0.53) |
| Switzerland | -1.01 | -1.22* | -1.20 |
|  | (0.64) | (0.73) | (1.01) |
| Business dummy | -0.33 | -0.02 | -0.07 |
|  | (0.74) | (0.58) | (0.74) |
| Constant | -5.70*** | -5.36*** | -4.30*** |
|  | (1.15) | (1.24) | (1.47) |
| AIC | 159.31 | 165.85 | 160.37 |
| BIC | 192.35 | 198.89 | 193.41 |

Looking at the biodiversity state variables that users most need and yet have low levels of access (as depicted in Figure 1), the GLMs show that there was no blockage significantly linked to organizations being unable to access data on habitat extent (Supplemental Materials S11), although blockages relating to accessing habitat quality data were significantly linked to data being unavailable (See Supplemental Materials S12; relevant data collection methods unknown or unavailable; β=-0.30, p<0.01).

Species population data was the second most needed type of data by biodiversity data users. The GLMs indicate that seven blockages were significantly linked to organizations being unable to access different types of species population data (Table 4).

**Table 4.** Generalized linear models for blockages against access to different types of species population data. Note: n = 93. DV = dependent variable. Robust standard errors in parentheses. Models where blockages are p<0.05 are presented here; full models available in Supplemental Materials. *** p<0.01, ** p<0.05, * p<0.1.

| Variables | DV: Access to species population data |  |  |  |  |  |  |  |  |
| --- | --- | --- | --- | --- | --- | --- | --- | --- | --- |
|  | Presence |  |  | Abundance |  | Range |  |  |  |
| Blockages |  |  |  |  |  |  |  |  |  |
| No monitoring and evaluation systems in place | -0.17** |  |  |  |  | -0.18** |  |  |  |
|  | (0.09) |  |  |  |  | (0.08) |  |  |  |
| Lack of technical capacity for data collection, analysis, and use |  | -0.18** |  |  |  |  | -0.21** |  |  |
|  |  | (0.09) |  |  |  |  | (0.09) |  |  |
| Lack of funding for data collection, analysis, and use |  |  | -0.22*** |  |  |  |  |  |  |
|  |  |  | (0.08) |  |  |  |  |  |  |
| Unavailable: cannot access from other sources |  |  |  | -0.36** |  |  |  |  |  |
|  |  |  |  | (0.15) |  |  |  |  |  |
| Unavailable: relevant data collection methods unknown or unavailable |  |  |  |  | -0.35** |  |  |  |  |
|  |  |  |  |  | (0.14) |  |  |  |  |
| Lack of understanding of the importance of monitoring in my organization |  |  |  |  |  |  |  | -0.24** |  |
|  |  |  |  |  |  |  |  | (0.10) |  |
| Lack of consensus about what to monitor |  |  |  |  |  |  |  |  | -0.21** |
|  |  |  |  |  |  |  |  |  | (0.09) |
| Controls |  |  |  |  |  |  |  |  |  |
| Female | 0.20 | 0.24 | 0.21 | -0.07 | -0.12 | -0.03 | 0.01 | -0.11 | -0.05 |
|  | (0.29) | (0.29) | (0.30) | (0.30) | (0.30) | (0.26) | (0.27) | (0.27) | (0.27) |
| Prefer not to specify (Gender) | -1.20*** | -0.95*** | -1.09*** | -0.50 | -0.52 | -2.04*** | -1.74*** | -2.15*** | -1.99*** |
|  | (0.29) | (0.36) | (0.28) | (0.33) | (0.32) | (0.28) | (0.37) | (0.25) | (0.28) |
| Age: 25-34 | -1.08*** | -0.96*** | -0.70* | -0.71 | -0.81* | -1.86*** | -1.72*** | -1.82*** | -1.71*** |
|  | (0.34) | (0.36) | (0.36) | (0.46) | (0.46) | (0.36) | (0.38) | (0.36) | (0.38) |
|  | Presence |  |  | Abundance |  | Range |  |  |  |
| Age: 35-44 | -0.83*** | -0.82*** | -0.56*** | -0.64*** | -0.65*** | -1.82*** | -1.79*** | -1.76*** | -1.79*** |
|  | (0.09) | (0.09) | (0.15) | (0.15) | (0.14) | (0.08) | (0.09) | (0.10) | (0.09) |
| Age: 45-54 | -1.25*** | -1.27*** | -0.98*** | -0.99*** | -0.99*** | -1.98*** | -1.99*** | -1.92*** | -1.98*** |
|  | (0.30) | (0.30) | (0.31) | (0.31) | (0.31) | (0.29) | (0.29) | (0.30) | (0.28) |
| Age: 55-64 | -0.95*** | -0.92*** | -0.75*** | -0.86** | -0.70* | -1.99*** | -1.95*** | -2.08*** | -1.96*** |
|  | (0.29) | (0.30) | (0.27) | (0.36) | (0.36) | (0.30) | (0.31) | (0.29) | (0.28) |
| Age: 65+ | -0.90*** | -0.93*** | -0.01 | -0.10 | -0.13 | -1.66*** | -1.70*** | -1.74*** | -1.68*** |
|  | (0.23) | (0.23) | (0.33) | (0.29) | (0.28) | (0.26) | (0.25) | (0.25) | (0.27) |
| Ghana | 0.36 | 0.43 | 0.36 | 0.23 | 0.26 | 0.27 | 0.33 | 0.35 | 0.26 |
|  | (0.29) | (0.30) | (0.29) | (0.36) | (0.36) | (0.30) | (0.30) | (0.30) | (0.30) |
| Switzerland | 0.82*** | 0.85*** | 0.75** | 0.33 | 0.20 | 0.72** | 0.76*** | 0.67** | 0.60** |
|  | (0.31) | (0.32) | (0.32) | (0.35) | (0.37) | (0.28) | (0.29) | (0.28) | (0.28) |
| Business dummy | 0.07 | 0.08 | -0.06 | 0.23 | 0.12 | 0.23 | 0.24 | 0.23 | 0.23 |
|  | (0.30) | (0.29) | (0.30) | (0.30) | (0.30) | (0.30) | (0.29) | (0.28) | (0.30) |
| Constant | 4.71*** | 4.68*** | 4.75*** | 4.94*** | 4.93*** | 5.56*** | 5.57*** | 5.63*** | 5.63*** |
|  | (0.30) | (0.32) | (0.33) | (0.54) | (0.52) | (0.27) | (0.28) | (0.26) | (0.30) |
| AIC | 291.29 | 291.29 | 289.79 | 307.24 | 307.54 | 289.50 | 288.65 | 286.77 | 288.47 |
| BIC | 314.09 | 314.09 | 312.58 | 330.03 | 330.33 | 312.29 | 311.44 | 309.57 | 311.26 |

There were only a handful of other data types significantly affected by specific blockages (see Supplemental Materials). The most common blockage linked to specific data access was data being unavailable (either by lack of access from other sources or unknown collection methods), which affected access to data on agriculture or aquaculture (Supplemental Material S17) and ecosystems services (soil quality; Supplemental Material S26).

Generally, there were more differences between Switzerland and the other two countries. For example, Switzerland had significantly different scores (p<0.05) from the other countries when accessing data for species populations (Table 4), habitat quality (Supplemental Materials S12), and ecosystem services (Supplemental Materials S23-S25). Controls for gender indicated that there were no significant differences in access to most types of biodiversity data between men and women.

## DISCUSSION

This study demonstrated that, although biodiversity data needs and data access vary between data user groups and between countries, the data most sought after across stakeholders relate to habitat state and species populations. However, these types of data are also among the ones with the biggest gap between access and use. While several studies have explored data availability, few have compared access with defined needs. Moreno et al. (2023) surveyed data users in East Africa and compared their needs with the availability of data in global biodiversity databases. They found that difficulty in accessing data was positively correlated with the importance of the data for conservation projects. As with our study, data most often needed but most difficult to access included parameters around species populations and habitat state. Given that the attainment of conservation goals and the impact of interventions can only be verified by a measured improvement in the state of habitats and species (Stephenson 2019; Nature Positive Initiative 2026), the challenges faced in accessing such data by diverse stakeholders is a cause for concern. In addition to the gap in data on progress with conservation action, this may, at least in part, explain the weak reporting of biodiversity performance by several actors. A review of reports from plant and animal conservation projects (Badalotti et al. 2022) found that less than one fifth of projects provided data on species populations. Similarly, only between one third and one half of national reports to the Convention on Biological Diversity include adequate data (Bubb 2013, Koh et al. 2022) and, of US companies sampled, only 12% made biodiversity disclosures (Carvajal et al. 2022). Lack of adequate data collection and reporting are likely to perpetuate geographic and taxonomic biases and gaps in species knowledge and biodiversity data sets (Hughes et al. 2021, Lindken et al. 2024).

Factors affecting data access are complex and hard to discern, but our results suggest they vary between user groups. However, the most frequently cited data blockages across user groups in our three study countries related primarily to issues of resources (inadequate funding) and organizational capacity (a lack of monitoring systems, no knowledge of what indicators to measure or which data collection methods to use, and an inability to access secondary data). Issues of inadequate capacity and funding are often recognised as key blockages to data access and use (Hochkirch et al. 2020, Hughes et al. 2021) but this is one of the first studies to quantify the biggest blockages across data users in multiple countries and to link those blockages to data needs. Our findings reflect two of the three most common blockages to data access identified by stakeholders in East Africa which also relate to inadequate funding and lack of capacity to process and analyse data (Moreno et al. 2023). Many companies are also often not equipped with the necessary expertise to collect and interpret biodiversity data (Heikkinen et al. 2023).

But, while across regions there may be some common trends, our study also demonstrated national differences in both data needs and data access. In general, blockages for Swiss data users were significantly different from Colombia and Ghana when accessing data for species, habitats, and ecosystem services. This could be related to a number of socio-economic, biological and institutional factors, although there have been few direct comparisons of nations in terms of capacity for data collection. Countries with larger economies, as measured by Gross Domestic Product (GDP), tend to have more species monitoring programmes (Moussy et al. 2022) and more biodiversity observations generally (Hughes et al. 2021). However, while Switzerland has a GDP 2.4 times larger than Colombia and 11.5 times larger than Ghana (World Bank 2025), lack of funding for data collection was the second most common blockage identified by Swiss organizations in our survey. This suggests that data access is less correlated with resources per se and more associated with priorities and budgeting at the organizational level. The size of each country and the number of species and habitats it contains might also be expected to influence data access. For example, Switzerland has fewer species to monitor than the other two larger tropical countries studied, as reflected in the number of species that have been assessed for the IUCN Red List of Threatened Species (IUCN 2025): 1,949 in Switzerland (41,285 km^2^) compared with 4,354 in Ghana (238,533 km^2^) and 15,881 in Colombia (1,140,619 km^2^).

We found that data blockages significantly associated with a lack of primary data collection included an absence of organizational biodiversity goals and monitoring systems. This further supports the idea that “good planning is a prerequisite for good monitoring” (Stephenson 2019). But why are stakeholders struggling to access the data they need on habitats and species?

Problems accessing both habitat quality and species abundance data were significantly associated with relevant data collection methods being unknown or unavailable to the organization. In contrast, no specific blockages were associated with accessing data on habitat extent, and species presence (and the related metric of species range) was more affected by funding and capacity issues and the absence of monitoring systems. This may reflect the fact that, while national, regional and global databases may be able to provide data on habitat extent or species presence, habitat quality and the abundance of species are more complicated metrics that generally require direct, site-level measurement. Species abundance (which can also be used as a proxy for habitat quality) is particularly difficult and requires the use of appropriate tools, methods and protocols, with some modern techniques like passive acoustic monitoring, less able to provide accurate density measures (Zwerts et al. 2021).

Compared with other data users, we found that businesses are more likely to need access to data on threats, from habitat fragmentation to infrastructure, agriculture and mining. This may reflect the growing importance to the business sector of measuring and disclosing environmental risk and impacts; prior research in business management has highlighted that business leaders generally lack knowledge around risks that arise from biodiversity (Overbeek et al. 2013, Boiral et al. 2019).

Businesses are less likely to collect primary data or use secondary data than other groups, and this may be linked to the finding that they are significantly more likely to be unclear on what biodiversity indicators to use. The growing number of biodiversity metrics and tools specifically designed for business can be confusing (Stephenson & Walls 2022), as is the rapid development of environmental disclosure standards such as the European Sustainability Reporting Standards within the EU Corporate Sustainability Reporting Directive. A low and uneven uptake of monitoring methods and availability of data within business sectors “varies across regions and countries, reflecting differences in data availability and technical capacity” (IPBES, 2026). The problem is further compounded by the lack of consistency between business metrics and those metrics used by other actors. While the Kunming-Montreal Global Biodiversity Framework focuses its goals, targets, and indicators on species and habitats in terrestrial, marine, and freshwater ecosystems (CBD 2022), the Taskforce on Nature-Related Financial Disclosures framework (TNFD 2023) uses realms (land, freshwater, ocean atmosphere), biomes, environmental assets, and ecosystem services. The Science-Based Targets Network guidance for business separates biodiversity from other aspects of nature such as climate, freshwater, land, and ocean (SBTN 2024). It is therefore unsurprising that many businesses are left confused and, and a result, biodiversity reporting and disclosure are limited (Adler et al., 2018; Skouloudis et al., 2019; Azizi et al., 2025; Kashyap et al., 2025). While some companies may have strong commitments to conserving biodiversity, achieving and reporting on these commitments often remain limited (Serrentino et al. 2024).

Our work therefore underlines the importance of improving the connection and collaboration between businesses and the conservation science and practice community that has been planning and monitoring biodiversity for decades. Co-operation and partnerships are enshrined within the Convention on Biological Diversity (CBD 2022) and are also key enabling conditions for improving biodiversity data access and use (Stephenson et al. 2017). This includes encouraging actors from different sectors to work together, including for the development of more harmonized indicators (McKenzie et al. 2025) and appropriate science-policy platforms and fora (Young et al. 2014).

Best practices need to be applied in choosing and using indicators and monitoring methods. While numerous standards, guidelines and tools are available, they need to be synthesised and shared more widely to help the diverse data users identify which indicators and monitoring methods and tools to use. Initiatives to build on include efforts by the Global Biodiversity Information Facility to build capacity through networks of data holders and users (GBIF 2019), a toolbox developed by GEOBON (the Group on Earth Observations Biodiversity Observation Network) to support national observation networks (GEOBON 2025, Schmeller et al. 2017), and guidelines, tools and data sources shared by the International Union for Nature Conservation, IUCN (e.g., IUCN SSC Species Monitoring Specialist Group 2025a,b, Stephenson & Carbone 2021).

In conclusion, global ambitions to conserve and restore biodiversity and produce nature-positive outcomes will only be possible through results-based, data-driven planning and adaptive management so that we can replicate what works well and improve what does not. This will require a concerted effort from governments, civil society, companies and academia to invest in the necessary internal capacity and external partnerships to enhance data collection, sharing and use and unblock the flow of biodiversity data.

## Acknowledgements

We are grateful to all the people in Colombia, Ghana and Switzerland who took time to complete our survey and thereby contribute to this work. Participants at a project workshop in September 2023 provided useful ideas and feedback on some of the preliminary results. We would like to thank Lena Kolecek for her invaluable help with project administration. The project was funded by a grant from the Swiss Network for International Studies (SNIS) to the University of Lausanne.

## Data availability

Data analyses are shared in Supplementary Materials; additional data are available from the authors on request.

## Ethical standards

The study, its empirical methodology and implementation were approved as ethically acceptable and in accordance with the regulations of the Ethics Committee of the University of St. Gallen under reference number HSG-EC-20240227.

## Author Contribution Statement

PJS conceived the idea; PJS, LF, JLW, MCLM & YN-B designed the project and methodology; KMMU, JAAM & LS collected and analysed the data; KMMU conducted formal statistical analyses; PJS and KMMU led the writing of the manuscript; all authors contributed critically to the manuscript drafts and gave final approval for publication.

